# Hexavalent Sperm-Binding IgG Antibody Released from Self-Dissolving Vaginal Film Enables Potent, On-Demand Non-Hormonal Female Contraception

**DOI:** 10.1101/2021.04.19.440503

**Authors:** Bhawana Shrestha, Kathleen Vincent, Alison Schaefer, Yong Zhu, Gracie Vargas, Massoud Motamedi, Kelsi Swope, Josh Morton, Carrie Simpson, Henry Pham, Miles B. Brennan, Michael H. Pauly, Larry Zeitlin, Barry Bratcher, Kevin J. Whaley, Thomas R. Moench, Samuel K. Lai

## Abstract

Non-hormonal products for on-demand contraception are a global health technology gap, motivating us to pursue the use of sperm-binding monoclonal antibodies as a strategy to enable safe, effective, desirable, on-demand contraception. Here, using cGMP-compliant Nicotiana-expression system, we produce an ultra-potent sperm-binding IgG antibody possessing 6 Fab arms per molecule that bind a well-established contraceptive antigen target, CD52g. We term this hexavalent antibody “Fab-IgG-Fab” (FIF) to reflect its molecular orientation. The Nicotiana-produced FIF exhibits at least 10-fold greater sperm agglutination potency and kinetics than the parent IgG, while preserving Fc-mediated trapping of individual spermatozoa in mucus. We formulate the Nicotiana-produced FIF into a polyvinyl alcohol-based water-soluble contraceptive film, and evaluate its potency in reducing progressively motile sperm in the sheep vagina. Two minutes after vaginal instillation of human semen, no progressively motile sperm are recovered from the vaginas of sheep receiving FIF-Film. In contrast, high numbers of progressively motile sperm are recovered from sheep receiving a placebo film control. Our work supports the potential of highly multivalent contraceptive antibodies to provide safe, effective, on-demand non-hormonal contraception.

## 1. Introduction

Despite the availability of potent and low-cost, long-acting, reversible contraceptives (LARCs), many women continue to use on-demand contraceptives due to infrequent sexual activity. In addition, many women strongly prefer non-hormonal contraceptives because of the real and/or perceived side-effects associated with existing hormonal methods.^[1–3]^ Indeed, the FDA-approved Vaginal Contraceptive Film (VCF) meets the contraceptive needs of many women as it provides a contraceptive method that is women-controlled, inexpensive, non-hormonal, discrete, and readily available over the counter. Unfortunately, VCF and most other spermicides use nonoxynol-9 (N9) as an active ingredient. N9 can damage the mucosal surfaces by disrupting the vulvar, vaginal, and cervical epithelium, and substantially increases the risks of sexually transmitted infections.^[4–6]^ We believe there is a substantial unmet need for alternatives that can offer effective on-demand contraception, and are free of exogenous hormones or detergents.

Anti-sperm antibodies (ASA) to surface antigens on sperm represent a promising class of molecules that could enable safe, on-demand, non-hormonal contraception. ASAs found in the vaginal secretions of some immune infertile women could prevent fertilization by stopping sperm from reaching the egg via two distinct mechanisms.^[7, 8]^ First, ASAs can agglutinate multiple motile sperm into clumps that stop forward progression.^[9,10]^ This mechanism is most effective at high sperm concentrations, and is more potent with polyvalent antibodies (Abs) such as IgM. Second, ASAs can trap individual spermatozoa in mucus by forming multiple low-affinity Fc-mucin bonds between sperm-bound ASA and mucin fibers, resulting in individual sperm that simply shake in place, unable to assume progressive motility needed to reach the upper reproductive tract.^[11]^ Over time, sperm that are agglutinated or immobilized in mucus either die or are eliminated from the female reproductive tract by natural mucus clearance mechanisms.

Years ago, the discovery of the contraceptive potential of ASAs motivated the development of contraceptive vaccines. ASAs elicited by vaccination with sperm antigens offered considerable contraceptive efficacy, but this approach stalled due to unresolved variability in the intensity and duration of the vaccine responses in humans, as well as concerns that active vaccination might lead to irreversible infertility.^[12–14]^ In contrast, topical delivery of pharmacologically active doses of ASA in the vagina can overcome many of the key drawbacks of contraceptive vaccines by providing consistent amounts of antibodies needed without risks of inducing immunity to sperm, thus making possible both consistently effective contraception and rapid reversibility. In good agreement with this concept, vaginal delivery of a highly multivalent anti-sperm IgM reduced embryo formation by 95% in a highly fertile rabbit model.^[15]^

This approach of topical passive immunocontraception has not been reported in humans, due in part to manufacturing and purification challenges with polyvalent Abs such as sIgA and IgM, and the lower agglutinating potencies of IgG. To overcome these challenges, we report here a highly multivalent IgG that possesses 6 Fabs per IgG molecule, with Fab domains interspersed by flexible glycine-serine linkers arranged in a **F**ab-**I**gG-**F**ab orientation; we term this molecule FIF **(Figure 1A).** To determine whether FIF may be useful for on-demand contraception, we produced FIF using a cGMP-compliant *Nicotiana benthamiana* manufacturing platform, and formulated the FIF into a dissolvable vaginal film comprised of polyvinyl alcohol. We report here the *in vitro* characterization and *in vivo* potency of this novel FIF Film.

**Figure 1.**
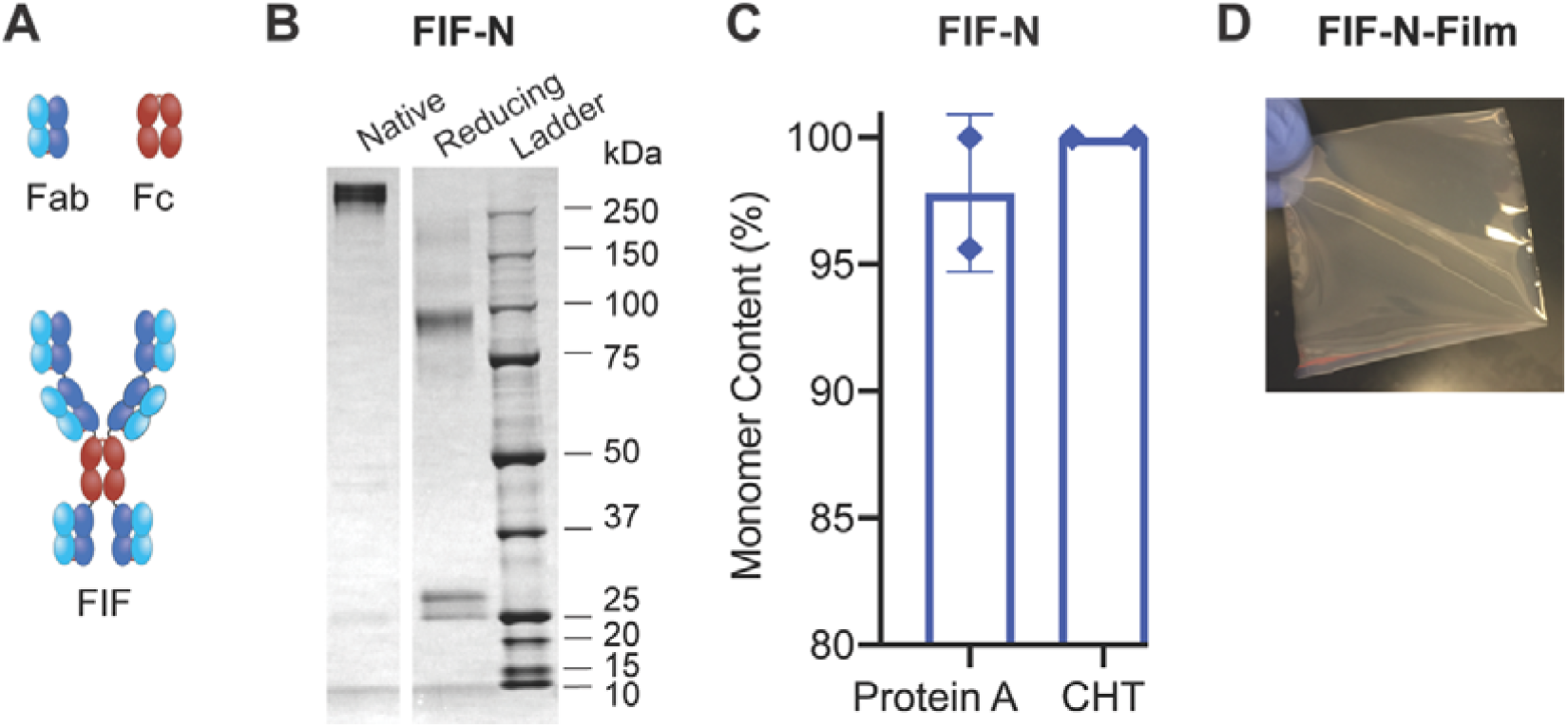
Production of FIF-N-Film. **(A)** Schematic diagrams of anti-sperm Fab-IgG-Fab (FIF). The additional Fab is linked to the N-terminal and C-terminal of parent IgG using flexible glycine-serine linkers to assemble FIF. **(B)** SDS-PAGE analysis of FIF-N in native (non-reducing) and reducing conditions. Non-reducing SDS-PAGE showcases the total molecular weight of the Ab and reducing SDS-PAGE displays the molecular weight of the individual heavy chain and light chain of Ab. **(C)** Demonstration of the homogeneity of FIF-N after protein A and ceramic hydroxyapatite (CHT) Chromatography using high performance liquid size exclusion chromatography (HPLC-SEC) analysis. Y-axis indicates the total percentage of Abs representing their theoretical molecular weights. **(D)** Image of watersoluble polyvinyl alcohol (PVA) film comprising of *Nicotiana-produced* FIF Ab.

## 2. Results

### 2.1. cGMP production of FIF in *N. benthamiana*

Efficient agglutination requires ASA to bind a ubiquitous antigen that is highly expressed on the surface of human sperm. For these reasons, we chose to engineer a monoclonal antibody (mAb) targeting a unique glycoform of CD52 (hereafter referred to as CD52g) that was previously shown to be produced and secreted by epithelial cells lining the lumen of the epididymis, and present on sperm, white blood cells in semen, and the epithelium of the vas deferens and seminal vesicles.^[16,17]^ The CD52g glycan-based antigen appears to be universally present on all human sperm while absent in most other tissues.^[17]^ Using a Fab-domain isolated from a healthy but immune infertile woman, we designed a 6 Fab antibody construct, cloned the sequences into the magnICON^®^ vector system, and transfected *Nicotiana benthamiana* using agrobacterial-infiltration process.^[18–22]^ This system allows for rapid and scalable production of full-length mAbs in two weeks; the same system has been used to produce various cGMP-compliant mAbs for clinical studies.^[23]^ To generate mAbs with homogeneous mammalian glycans, we used a transgenic strain, Nb7KOΔXylT/FucT of *N. benthamiana* which yields mAb with predominantly G0 N-glycans. Without optimization, the production yields of the *Nicotiana-produced* FIF (FIF-N) post-protein A chromatography were approximately 29 mg kg^-1^ of plant tissue **(Figure S1A)**. The mAbs were further purified using ceramic hydroxyapatite chromatography prior to further biophysical characterization. SDS-PAGE analysis demonstrated the correct assembly of FIF-N at its theoretical molecular weight, ~350 kDa (**Figure 1B**). Purified FIF contained >99% monomeric form as determined by high performance liquid size exclusion chromatography analysis **(Figure 1C and Figure S1B).** FIF-N demonstrated excellent stability, with no appreciable aggregation or degradation upon storage at room temperature for 3 weeks and freezing at −70 °C **(Figure S1, B and C).**

### 2.2. Production of FIF-N-Film

Polyvinyl alcohol (PVA) is a polymer routinely used in biomedical applications. Low molecular weight PVA is widely used in female reproductive health products suitable for intravaginal administration, with no appreciable vaginal toxicity or irritation. Similar to prior work in formulating a vaginal film releasing both an anti-HIV (VRC01) and anti-HSV (HSV8) mAb evaluated in a Phase 1 trial, we prepared water-soluble PVA films comprised of PVA 8-88 (67 kDa) together with 10 mg of FIF-N, using an aqueous casting method. As a control, an IgG-N-Film with 20 mg of anti-CD52g IgG was also prepared.^[24]^ Both films were fabricated to 2”x2” in dimensions, clear in visual appearance with few bubbles present, homogeneous, and resistant to tear **(Figure 1D)**. Both films showed no significant levels of endotoxin, and no detectable bioburden (CFU mL^-1^), indicating efficient and aseptic removal of potential contaminants **(Table S1)**.

### 2.3. FIF-N-Film possesses superior agglutination potency

We next assessed the sperm-agglutinating potencies of dissolved IgG-N and FIF-N films. We focused on assessing the reduction in progressive motile (PM) fraction of sperm, since it is the PM sperm fractions that reach the uterus and penetrate the zona pellucida to fertilize the egg. We first assessed the agglutination potencies of FIF-N-Film vs. IgG-N-Film using a sperm escape assay with purified sperm. The sperm escape assay uses Computer Assisted Sperm Analysis (CASA) to quantify the number of PM sperm that escapes agglutination over 5 mins when mixed with specific mAbs at different mAb and sperm concentrations. We elected to first assess agglutination at a low concentration of 5 million PM sperm mL^-1^, the minimal PM sperm concentration in semen associated with fertility, which limits sperm collision frequency and making it more challenging to achieve rapid and complete agglutination. FIF-N-Film exhibited at least 16-fold greater agglutination potency than IgG-N-Film, defined here as the minimal mAb concentration at which PM sperm are reduced by >98%. The minimum concentration of IgG-N-Film needed was ~6.25 μg mL^-1^, whereas just 0.39 μg mL^-1^ of FIF-N-Film was sufficient **(Figure 2A).**

**Figure 2.**
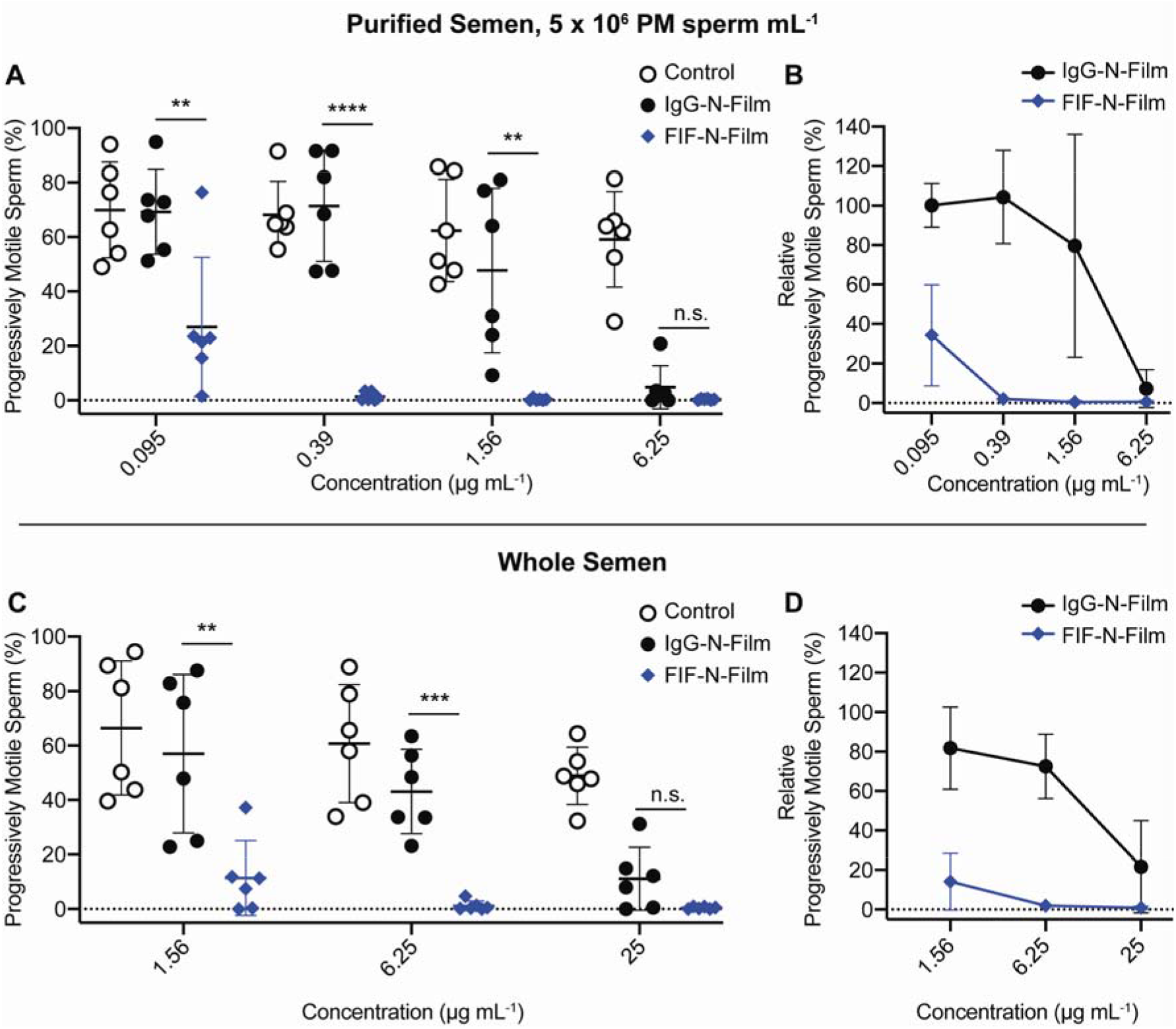
FIF-N-Film possesses markedly greater agglutination potency than IgG-N-Film. **(A)** Sperm agglutination potency of the IgG-N-Film and FIF-N-Film determined by quantifying PM sperm that escaped agglutination after Ab-treatment compared to pre-treatment condition using CASA. Purified sperm at the final concentration of 5 x 10^6^ PM sperm mL^-1^ was used. **(B)** Sperm agglutination potency of the Abs normalized to the media control. **(C)** Further assessment of sperm-agglutination potency of the IgG-N-Film and FIF-N-Film using whole semen. **(D)** Sperm-agglutination potency of the IgG-N-Film and FIF-N-Film against whole semen normalized to the sperm washing media control. Data were obtained from N□=□6 independent experiments with 6 unique semen specimens. Each experiment was performed in duplicates and averaged. P values were calculated using a oneway ANOVA with Dunnett’s multiple comparisons test. *P□<□0.05, **P□<□0.01, ***P < 0.001 and ****P < 0.0001. Data represent mean ± standard deviation.

To confirm efficient agglutination also occur with native semen, we further assessed the agglutination potency of the FIF-N-Film vs IgG-N-Film using whole semen. FIF-N-Film again exhibited at least 10-fold greater agglutination potency than IgG-N-Film **(Figure 2C)**. Both FIF-N-Film and IgG-N-Film required ~16-fold more mAb to achieve >98% agglutination of PM sperm in whole semen compared to in purified motile sperm, likely due to CD52g present on other components in whole semen, including non-PM sperm, seminal leukocytes, as well as on exosomes from the epithelium of the vas deferens and seminal plasma.^[25]^

### 2.4. FIF-N-Film exhibits faster sperm agglutination kinetics

For effective vaginal immunocontraception based on limiting sperm motility in mucus, mAbs must agglutinate/immobilize sperm before they reach the upper reproductive tract; thus, rapid reduction of PM sperm is likely an important factor in contraceptive efficacy. Thus, we next quantified the kinetics of sperm agglutination by quantifying the number of PM motile sperm present at 30 s intervals following treatment of purified sperm (5 million PM sperm mL-1) with IgG-N-Film and FIF-N-Film. IgG-N-Film reduced PM sperm by ≥90% within 90 s in 5 of 6 semen samples at 6.25 μg mL^-1^, but failed to do so in 6 of 6 samples at 1.56 μg mL^-1^ **(Figure 3A).** In contrast, FIF-N-Film agglutinated ≥90% of PM sperm within 30 s in all cases at both 6.25 μg mL^-1^ and 1.56 μg mL^-1^ concentrations **(Figure 3A)**. Even at 0.39 μg mL^-1^, FIF-N-Film still agglutinated ≥90% of PM sperm within 90 s in 5 of 6 samples. Notably, the agglutination kinetics of FIF-N-Film was markedly faster and more complete than the parent IgG at all mAb concentrations and across all time points **(Figure 3B)**.

**Figure 3.**
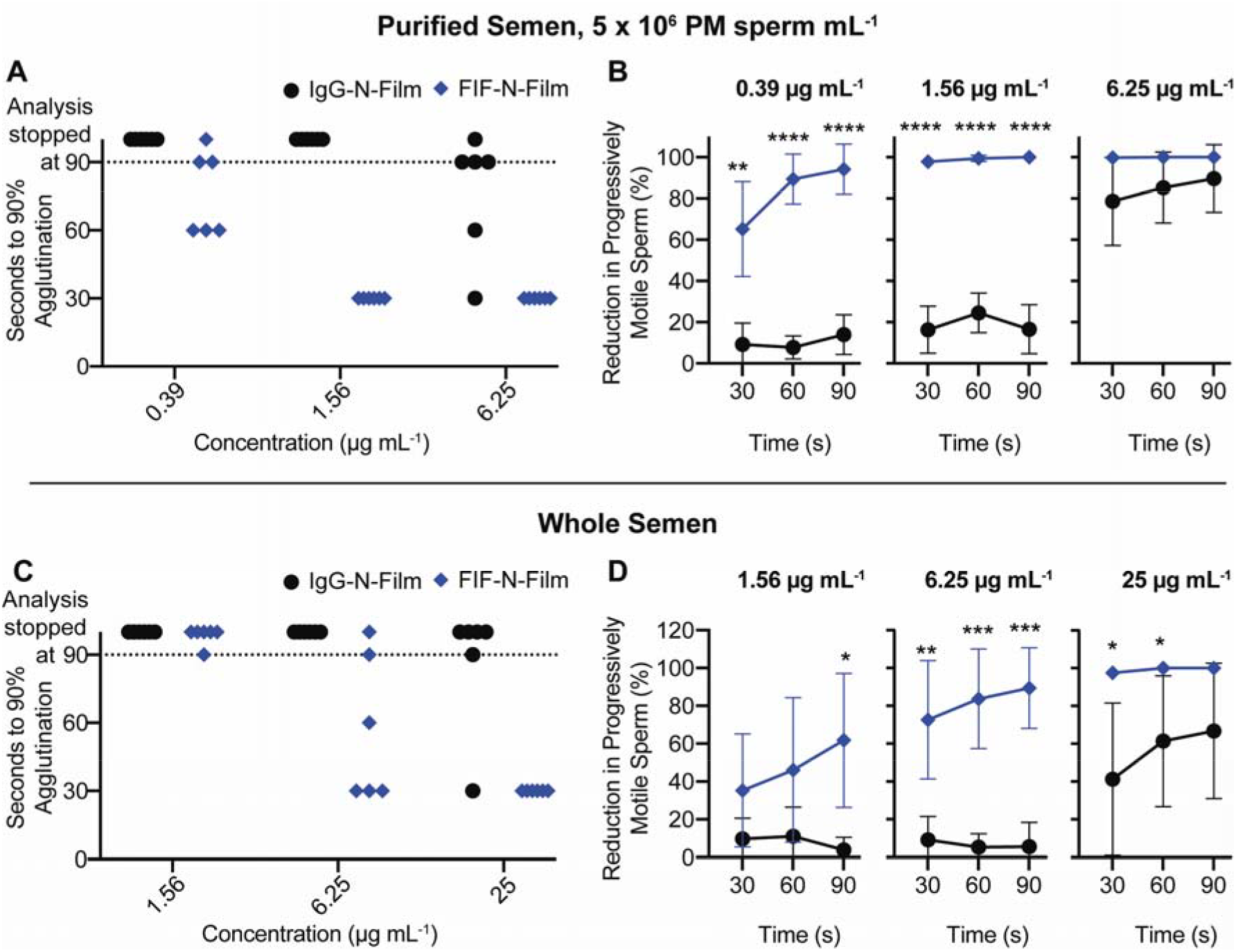
FIF-N-Film exhibits markedly faster agglutination kinetics than IgG-N-Film. **(A)** Sperm agglutination kinetics of IgG-N-Film and FIF-N-Film measured by quantifying the time required to achieve 90% agglutination of PM sperm compared to sperm washing media control. **(B)** The rate of sperm agglutination determined by measuring the reduction in the percentage of PM sperm at three timepoints after Ab-treatment compared to sperm washing media control. Purified sperm at the final concentration of 5 x 10^6^ PM sperm mL^-1^ was used. **(C)** Sperm agglutination kinetics and **(D)** The rate of sperm agglutination assessed for IgG-N-Film and FIF-N-Film using whole semen. Data were obtained from N□=□6 independent experiments with 6 unique semen specimens. Each experiment was performed in duplicates and averaged. P values were calculated using a one-tailed t-test. *P□<□0.05, **P□<□0.01, ***P < 0.001 and ****P < 0.0001. Data represent mean ± standard deviation.

Similar to the sperm escape assay, we also assessed agglutination kinetics of FIF-N-Film vs. IgG-N-Film using whole semen. Again, a higher concentration of FIF-N-Film and IgG-N-Film was required to obtain comparable agglutination kinetics vs. purified sperm. Nonetheless, FIF-N-Film exhibited markedly faster and more complete sperm agglutination kinetics than IgG-N-Film at all mAb concentrations and all time points in whole semen **(Figure 3D)**. At 25 μg mL^-1^, FIF-N-Film agglutinated ≥90% of PM sperm within 30 s in 6 of 6 whole semen samples while IgG-N-Film agglutinated ≥90% of PM sperm in 90 s in only 2 of 6 specimens at the same concentration **(Figure 3C)**. Lower sperm concentration (as found in semen from oligospermia, sub-fertile individuals) may limit sperm agglutination due to reduced likelihood of a sperm-sperm collision, whereas higher sperm amounts may saturate the agglutination potential. We thus further assessed whether FIF-N-Film can effectively reduce PM sperm at 1 million PM sperm mL^-1^ and 25 million PM sperm mL^-1^. FIF-N-Film maintained similar superior agglutination kinetics over IgG-N-Film across both lower and higher sperm concentrations **(Figure S2)**. These results underscore the increased potency for FIF-N-Film compared to the IgG-N-Film across diverse conditions.

### 2.5. FIF-N and FIF-Expi293 exhibits equivalent agglutination

To confirm that the production of FIF in *N. benthamiana* and their subsequent formulation into PVA films did not reduce their agglutination activity, we further compared the sperm agglutination potencies of FIF-N, before and after film formulation, to Expi293-produced FIF. At 0.39 μg mL^-1^, FIF-Expi293, FIF-N, and FIF-N from four dissolved FIF-N-Films all demonstrated comparable sperm agglutination potencies **(Figure S3A)**. Similarly, FIF-Expi293, FIF-N, and FIF-N from dissolved FIF-N-Films all agglutinated all sperm within 60 s in 3 of 3 samples at 1.56 μg mL^-1^ **(Figure S3B).** The agglutination kinetics profile of Expi293- and *Nicotiana-* produced FIF, pre- and post-film formulation, were also virtually identical **(Figure S3C).** These results underscore that neither production of FIF in *Nicotiana* nor formulation of FIF-N into films had any significant impact on the actual agglutination potencies of FIF.

### 2.6. FIF-N-Film traps individual spermatozoa in vaginal mucus

Previous work has shown that IgG and IgM Abs can retard the active motility of individual spermatozoa in mucus despite continued vigorous beating action of the sperm flagellum; clinically, this is referred to as the “shaking phenomenon”.^[11]^ This muco-trapping function is similar to recent observations with Herpes Simplex Virus (HSV), whereby multiple HSV-bound IgGs formed polyvalent adhesive interactions between their Fc domains and mucin fibers in cervicovaginal mucus (CVM).^[26,27]^ Anti-HSV IgG-mediated effective trapping of individual viral particles in CVM, and blocked vaginal Herpes transmission in mice.^[26]^ We thus assessed whether FIF-N-Film can reduce progressive motility of fluorescently labeled spermatozoa in the relatively thin (low viscosity) CVM using multiple particle tracking. FIF-N-Film reduced progressively motile spermatozoa to the same extent as the IgG-N-Film, indicating that the addition of Fabs to both the N- and C-terminus of the IgG did not interfere with Fc-mucin crosslinking **(Figure 4).**

**Fig. 4.**
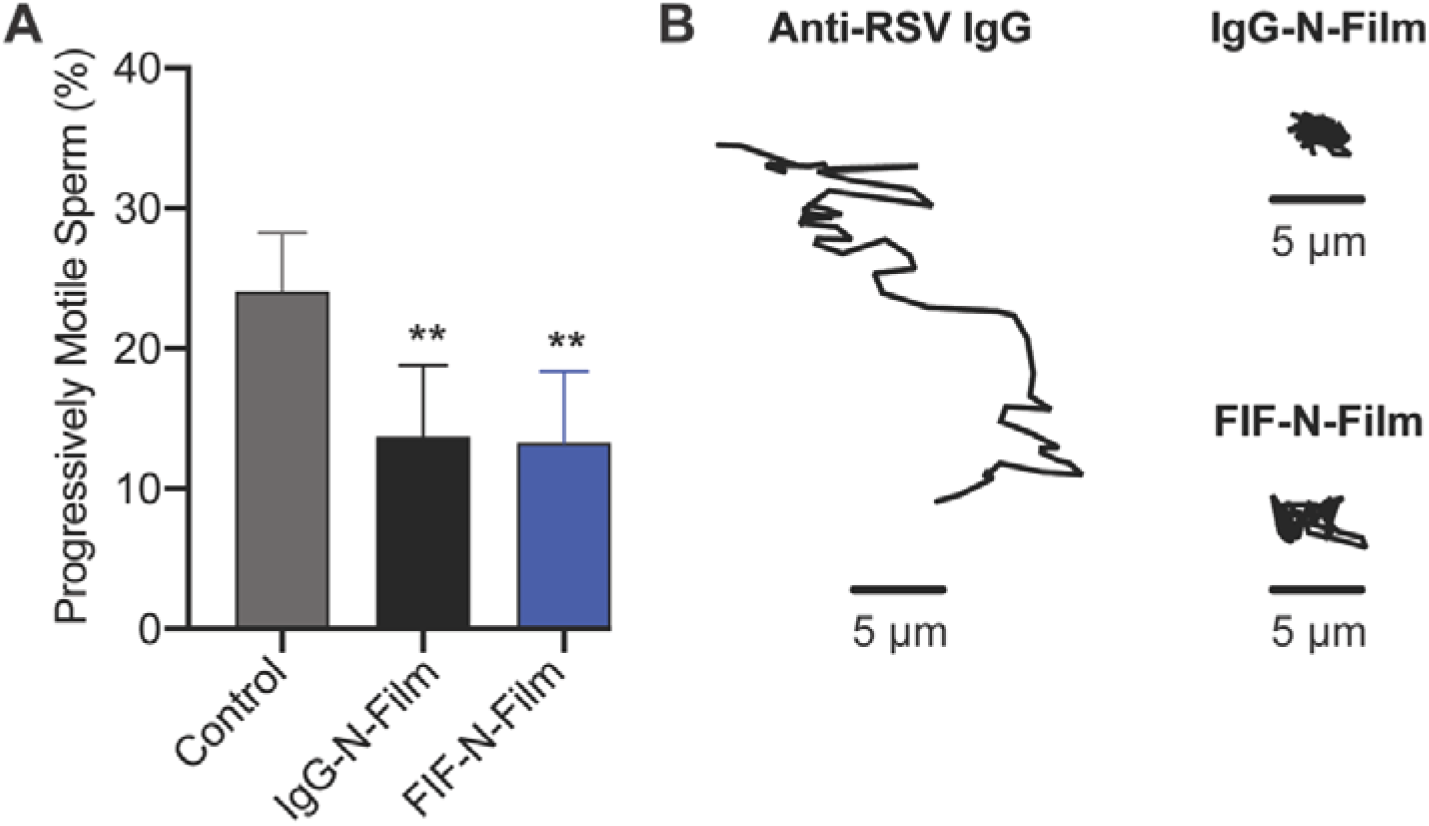
FIF-N-Film maintains the trapping potency of IgG-N-Film. **(A)** The trapping potency of the indicated Abs (25 μg mL-1) measured by quantifying fluorescently labeled PM sperm in Ab-treated CVM using neural network tracker analysis software. Purified sperm at the final concentration of 5.8 x 10^4^ PM sperm mL-1 was used. Data were obtained from N□=□6 independent experiments with 6 unique combinations of semen and CVM specimens. P values were calculated using a one-tailed t-test. *P□<□0.05 and **P□<□0.01. Data represent mean ± standard deviation. **(B)** Representative 4 s traces of sperm within one standard error mean of average path velocity at a timescale τ of 1 □s in CVM treated with control (anti-RSV IgG), IgG-N-Film, and FIF-N-Film.

### 2.7. FIF-N-Film rapidly eliminates PM sperm in sheep vagina

Since the unique glycoform of CD52g is only found in human and chimpanzee sperm, there is no practical animal model to perform mating-based contraceptive efficacy studies.^[28]^ Instead, we designed a sheep study that parallels the human post-coital test (PCT), which assesses the reduction of PM sperm in the female reproductive tract (FRT) given that PM sperm are required for fertilization.^[29–33]^ Clinical PCT studies have proven to be highly predictive of contraceptive efficacy in clinical trials.^[30,34-39]^ The sheep vagina is physiologically and anatomically very similar to the human vagina, making it the gold standard for assessing vaginal products.^[40,41]^ We instilled either Placebo-Film (no mAb) or FIF-N-Film into the sheep vagina, allowed 30 mins for the film to dissolve, followed by brief simulated intercourse with a vaginal dilator (15 strokes), vaginal instillation of fresh whole human semen, brief simulated intercourse (5 strokes), and finally, recovery of the semen mixture from the sheep vagina 2 mins post semen instillation for immediate visual assessment of sperm motility via quantifying progressively motile sperm. Despite this exceptionally stringent criteria, FIF-N-Film reduced 100% of PM sperm in all four of the animals studied over two independent studies, with no observable PM sperm (**Figure 5**; p<0.0001). In contrast, there were high PM sperm fractions recovered from all four sheep receiving the placebo film, with a few to several hundred PM sperm counts in the microscopy field, comparable to those from sheep treated with saline control.

**Figure 5.**
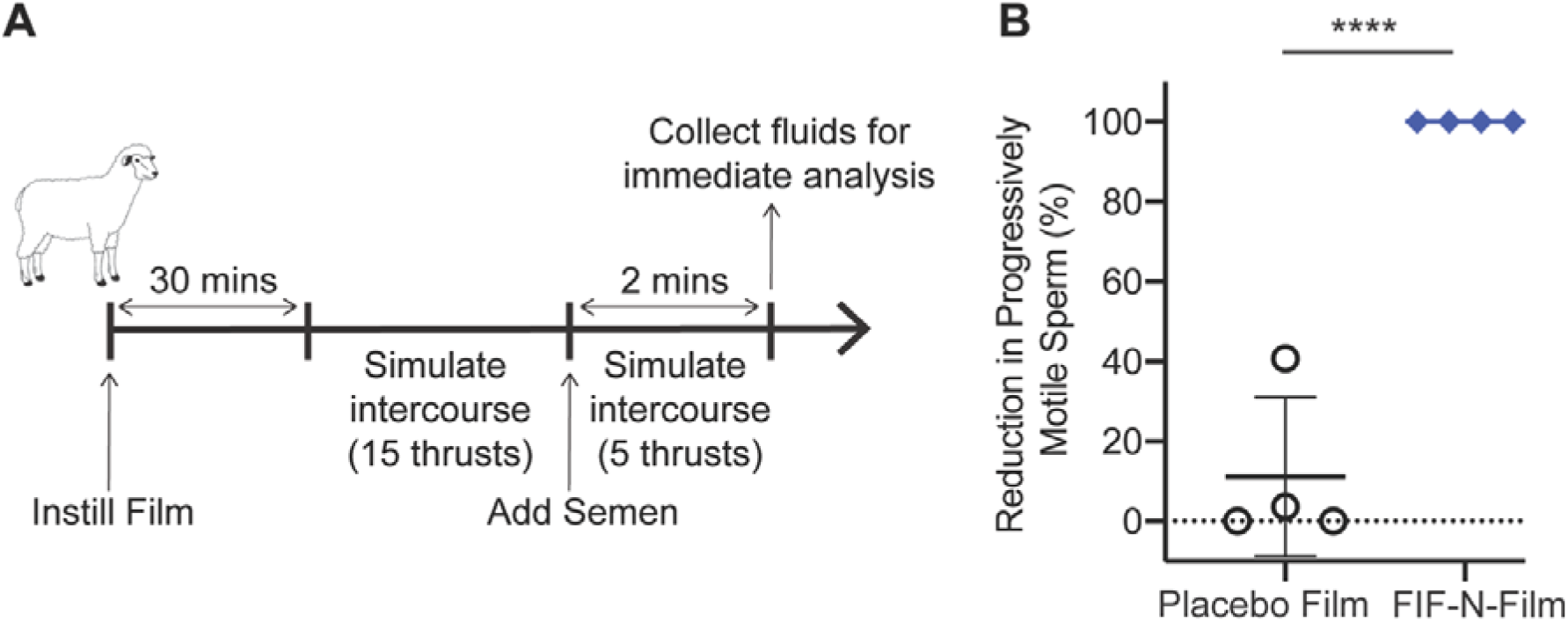
FIF-N-Film exhibits complete agglutination in surrogate sheep studies. **(A)** Schematic of the study design. **(B)** The potency of Placebo-Film and FIF-N-Film measured by quantifying PM sperm in sheep’s vaginal fluid after Ab- or Placebo-treatment compared to saline-treatment. Treatment administration was blinded, and quantifications were performed using a neural network tracker coded with sperm motility parameters. Data were obtained from N□=□3 independent experiments using 5 sheep per experiment. P values were calculated using a one-tailed t-test. *P□<□0.05, **P□<□0.01, ***P < 0.001 and ****P < 0.0001. Data represent mean ± standard deviation.

## 3. Discussion and Conclusion

The sperm agglutination potency of FIF-N-Film in sheep reported here is likely attributed in large part to the additional Fab arms of the FIF molecule. By delivering FIF directly to where it is needed i.e. the vagina, the fraction of FIF available to bind sperm is maximized, thereby enabling complete agglutination and immobilization of progressively motile sperm within just two minutes of semen exposure. In contrast, only a tiny fraction of systemically delivered mAb will be available to bind sperm because of the large blood volume (~5L), distribution to non-target tissues, natural catabolic degradation, and finally limited and delayed distribution into the FRT, including the vagina. As a result, markedly lower total amount of FIF is needed with vaginal delivery to achieve contraceptive levels in the FRT compared to delivering the same mAb systemically. An added advantage of vaginal delivery is that the entire dose of FIF delivered is quickly available, without any delays in reaching C_max_ in the vagina from delayed extravasation from the systemic circulation. Vaginal IgG has a half-life of ~9 hrs; thus, even after 24 hrs, there will likely be sufficient quantities of FIF from the original 10 mg film to maintain effective sperm agglutination, given in vitro measurements that showed highly effective sperm agglutination even at FIF concentrations as low as ~390 ng mL^-1^.^[42]^

Decades ago, the high costs of mAb production and emphasis on systemic administration critically limited the feasibility of passive immunization with ASA as a strategy for non-hormonal contraception. However, the cost of mAb production has declined over the years due to advances in CHO cell production. It reportedly costs between $95-$200 per gram to produce currently marketed mAbs.^[43]^ If a CHO facility uses a continuous bioprocess system integrated with single-use bioreactors it has been predicted to reduce mAb manufacturing costs per gram to less than $15 per gram, or $3 for an average 200 mg dose of most systemic mAbs. Based on 10 mg of FIF per film that resulted in exceptional sperm agglutination potency in the sheep vagina, the costs to produce the needed amount of FIF per film in CHO is likely much less than $1. Since FIF exhibits considerable agglutination potencies down to 390 ng mL^-1^, additional dose optimization may further reduce the amount of FIF needed per film, thus further decreasing costs and improving scale.

mAbs based topical contraceptives, such as the FIF-N-film reported here, are likely to be safe due to their binding specificity, particularly when targeted to epitopes present primarily on sperm. Vaginally dosed mAbs are poorly absorbed into the systemic circulation, and the vaginal immune response is limited even when vaginally vaccinating with the aid of highly immunostimulatory adjuvants.^[44–46]^ Vaginal secretions naturally contain high levels of endogenous IgG (*i.e.* 1-2 mg mL^-1^, making it unlikely that vaginal delivery of FIF, which is comprised of fully human Fabs and Fc, would trigger inflammation, sensitization, or other local toxicities. Finally, PVA (67 kDa) film, which is widely used in pharmaceutical applications as well as in contraceptive products such as VCF, has been found to be safe and non-immunogenic to use.^[47,48]^ Altogether, these features make PVA film delivering mAb vaginally for immunocontraception likely to be exceptionally safe.

Typically, only ~1% of the ejaculated sperm enter the cervix, even fewer reaching the uterus, and only dozens of sperm (out of the ~200 million in the ejaculate) reach the neighborhood of the egg.^[49]^ Accordingly, poor sperm motility in mid-cycle cervical mucus and low total sperm count are considered good correlates to low conception rates. Human semen averages between 45-65 million sperm mL^-1^, 15 million sperm mL^-1^ marks the lowest 5th percentile in men with proven fertility, and <5 million sperm mL^-1^ is often considered severe oligospermia that correlates with very low fertility.^[50–52]^ These observations suggest a marked reduction of progressive sperm motility, even if incomplete (e.g. 10-fold reduction in PM sperm fractions), may likely provide substantial contraceptive efficacy. This expectation is also consistent with the observations that even under ideal circumstances, with unprotected intercourse on the cycle day of maximum fertility, the odds of conceiving are only about ~10%.^[53]^ This indicates that only a small (*i.e.* limiting) number of motile sperm would reach the egg per intercourse; thus, reducing progressive sperm motility in the vagina and cervical canal should proportionally reduce the likelihood of conceiving. These findings, together with the contraceptive success with topical ASA against rabbit sperm, suggest arresting progressive sperm motility in mucus using mAb (which can reduce PM sperm by >99.9%) should provide an effective form of contraception.^[15]^

One potentially important mechanism of vaginal HIV transmission is cell-associated HIV transmission, whereby HIV in immune cells of HIV+ semen facilitates direct cell-to-cell spread of the virus to target cells in the female reproductive tract.^[54]^ Cell-associated HIV transmission may be more efficient than cell-free HIV transmission, since intracellular viruses are not exposed to the same host restriction factors and innate immune molecules the FRT. Since CD52g is adsorbed on the surface of immune cells originating from the male reproductive tract, it is possible that FIF can also agglutinate such immune cells and limit their access to target cells in the FRT, thereby limiting cell-associated vaginal HIV transmission. Combining contraception with the prevention of sexually transmitted infections is also an attractive public health strategy. With further reduction in manufacturing costs and greater availability of multi-metric ton manufacturing capacity for mAbs, it may be possible to create a cost-effective, on-demand multi-purpose technology product based on a cocktail of antiviral and anti-sperm mAbs that can simultaneously afford potent contraception and effective protection against STI transmission.

Polymeric vaginal films are advantageous for delivering active pharmaceutical ingredients (API) and preferred over other delivery methods due to enhanced bio-adhesive properties, ease of use, compact size and negligible vaginal leakage.^[55–59]^ Currently, multiple vaginal films with anti-retroviral microbicides are under development and evaluation.^[60-63]^ In a recent Phase I study, a vaginal film formulated with the microbicide drug candidate, dapivirine, was found to be safe and acceptable with uniform vaginal distribution while exhibiting considerable efficacy against *ex vivo* HIV-1 challenge model.^[57]^ Similarly, vaginal films could be formulated with contraceptive mAbs and microbicides or anti-fungal agents to achieve multipurpose prevention. Finally, it may be possible to formulate vaginal films to provide sustained release in the vagina spanning days to weeks.^[64,65]^

There are a number of limitations to our current study. First, we did not directly demonstrate efficacy by preventing pregnancies. We are unable to do so due to the unique antigen (CD52g) that our antibody targets: prior work has shown that, besides humans, only chimpanzees possess CD52g, and it is not possible to conduct chimpanzee studies in the U.S.^[28]^ Instead, for our in vivo proof-of-concept study, we were forced to adopt a sheep model designed to closely mimic the human post-coital test that is routinely used to assess the efficacy of sperm-targeted contraceptives in early phase clinical studies. Fortunately, the human cost-coital test has shown to correlate well with eventual efficacy in preventing pregnancies. Second, the precise dose of antibodies needed to ensure highly effective sperm agglutination remains not well understood. In the current study, to ensure success, we incorporated a relatively large dose of mAb (10 mg) into the vaginal film formulation. Although we expect this dose of mAb to be commercially viable (~$1 per film based on bulk manufacturing costs of mAb at $100 per gram), it is likely we can achieve effective agglutination of sperm with even lower quantities of mAb released from the film, which would translate to even lower costs. We are also pursuing the development of other vaginal delivery formats, such as an intravaginal ring that can afford sustained release of our mAbs across the potential fertility window, which may further reduce the dose needed.

## 4. Experimental Section

### 4.1. Experimental design and ethics

The objective was to assess the sperm-agglutinating and -trapping potency of PVA film formulated with *Nicotiana*-produced FIF Ab in vitro and in vivo. The in vitro studies using human semen and human cervicovaginal mucus samples were approved by the Institutional Review Board (IRB) of the University of North Carolina at Chapel Hill (IRB-101817). Prior to the collection of semen and mucus samples, informed written consents were obtained from all male and female subjects. Mass student emails and printed posters were utilized to recruit subjects for the UNC-Chapel Hill studies. The sheep surrogate post-coital test using human semen samples was approved by the IRB of the University of Texas Medical Branch (UTMB; IRB-180254). Informed written consent was obtained from the pre-screened male volunteers. Sheep studies were approved by the UTMB Institutional Animal Care and Use Committee (IACUC-0608038D) and utilized 5 female Merino crossbred sheep. IgG-N-Film and FIF-N-Film were dissolved in ultra-pure water before all in vitro experiments.

### 4.2. Construction of N. benthamiana expression vectors

The variable light (V_L_) and variable heavy (V_H_) DNA sequences for anti-sperm IgG antibody were obtained from the published sequence of H6-3C4 mAb.^[18,19]^ For the construction of expression vector encoding light chain (LC), a gene fragment consisting of V_L_ and C_λ_ DNA sequences was codon-optimized and synthesized using GeneArt^®^ gene synthesis services (ThermoFisher Scientific) and cloned into PVX viral backbone (Icon Genetics).^[20]^ For the construction of an expression vector containing IgG1 heavy chain (HC), a gene fragment consisting of V_H_ and C_H_1-C_H_2-C_H_3 DNA sequences was codon-optimized and synthesized using GeneArt^®^ gene synthesis services (ThermoFisher Scientific) and cloned into TMV viral backbone (Icon Genetics).^[20]^ For the construction of expression vector containing FIF HC, a gene fragment consisting of V_H_/C_H_1-(G_4_S)_6_ Linker-V_H_/C_H_1-C_H_2-C_H_3-(G_4_S)_6_ Linker-V_H_/C_H_1 DNA sequences was synthesized using GeneArt^®^ gene synthesis services (ThermoFisher Scientific) and cloned into TMV viral backbone (Icon Genetics).

### 4.3. Production of mAbs in Nb7KOΔXylT/FucT N. benthamiana

Briefly, IgG and FIF mAbs were expressed in *N. benthamiana* plants using “magnifection” procedure.^[23]^ Cloned expression vectors i.e. PVX-LC, TMV-IgG-HC, and TMV-FIF-HC were transformed into *Agrobacterium tumefaciens* strain ICF320 (Icon Genetics) and grown overnight at 28.0 °C followed by 1:000 dilution in infiltration buffer [10 mm MES (pH 5.5) and 10 mm MgSO_4_]. The combinations of diluted bacterial cultures (TMV-IgG-HC + PVX-LC and TMV-FIF-HC + PVX-LC) were used to transfect 4 wks old *N. benthamiana* plants (ΔXTFT glycosylation mutants) using vacuum infiltration. Using a custom-built vacuum chamber (Kentucky Bioprocessing), the aerial parts of entire plants were dipped upside down into the bacterial/buffer solution and a vacuum of 24’’ mercury was applied for 2 mins. Infiltrated plants were allowed to recover and left in the growth room for transient expression of antibodies. 7 days after infiltration, plants were harvested and homogenized in extraction buffer containing 100 mm Glycine, 40 mm Ascorbic Acid, 1m EDTA (pH 9.5) in a 0.5:1 buffer (L) to harvested plants (kg) ratio. The resulting green juice was clarified by filtration through four layers of cheesecloth followed by centrifugation at 10,000 g for 20 mins. Next, mAbs were captured from the clarified green juice using MabSelect SuRe Protein A columns (GE Healthcare). Protein A Columns were equilibrated and washed with buffer containing 50 mm Tris, pH 7.4, and bound protein were eluted with buffer containing 100 mm Acetic acid, pH 3. The eluates were immediately neutralized using 1 m Tris, pH 8.0. The eluted mAbs were further purified using equilibrated Capto Q columns (GE Healthcare) and flow-through fractions, which contain mAbs, were collected. The mAb-containing fractions were finally polished with CHT chromatography with type II resin (Bio-Rad). The CHT columns were equilibrated and washed with phosphate running buffer and eluted with running buffer containing NaCl.

### 4.4. Biophysical characterization of mAbs

SDS-PAGE at reducing and non-reducing conditions was performed to determine the molecular weight of FIF-N. Briefly, mAb (1 μg) was denatured at 70 °C for 10 min. Next, 0.5 m tris (2-carboxyethyl) phosphine (TCEP; 0.3 μL) was added as a reducing agent to the denatured protein for a reduced sample and incubated at room temperature for 5 min. After the incubation, samples were loaded, and the gel was run for 40 min at a constant voltage of 200 V. Bio-Rad Precision Protein Plus Unstained Standard was used as a protein ladder. Imperial Protein Stain (Thermo Scientific) was used to visualize the protein bands. The brightness and contrasts of the SDS-PAGE image were linearly adjusted using Image J software (Fiji).

HPLC-SEC was performed to determine the purity of IgG-N and FIF-N mAbs. The HPLC-SEC system consisted of a TSK Gel Super SW3000 column (Tosoh Biosciences) connected to Agilent 1260 HPLC system and a UV detector. The flow rate was maintained at 0.2 mL min^-1^. The column was equilibrated with 0.1 m sodium phosphate, 0.15 m NaCl buffer, pH 7.2 before loading the samples. Each mAbs (100 μg) were injected onto the column, and data were collected and analyzed using the ChemStation chromatography data system and software (Agilent). The proportion of monomers, aggregates, and fragments present in each mAb sample were calculated using ChemStation software (Agilent).

Endotoxin levels in mAbs were measured with Endosafe PTS (Charles River), which detects by measuring color-intensity related to endotoxin concentration.

Bioburden was determined for IgG-N and FIF-N by counting the number of colony-forming units that formed after mAbs were incubated overnight on the bacterial agar plate at 37 °C.

### 4.5. Production of IgG-N and FIF-N films

Films were manufactured using the solvent casting method^[55]^. Briefly, PVA 8-88 (67 kDa; 25%, w/w) was dissolved in MilliQ water. Next, IgG and FIF mAbs suspended in 10 mM Histidine + 0.005% Polysorbate 20, pH 6.5 were slowly added into the PVA solution followed by 200 mg mL^-1^ maltitol. The solution was stirred over 15 minutes to ensure uniform distribution of mAbs and to remove the entrapped air bubbles. The final uniform polymer solution was cast onto a polyester substrate attached to a glass plate using a 2”x2”x0.020” die press. The film sheet was allowed to dry for 20 min before it was removed from the substrate, and then cut into 2”x1.8” individual unit doses using a scalpel. Placebo film was prepared using the same method as described above except without drug substances in the polymer solution. IgG-N-Film and FIF-N-Film were dissolved in ultra-pure water prior to in vitro experiments.

### 4.6. Semen collection and isolation of purified motile sperm

Healthy male subjects were asked to refrain from sexual activity for at least 24 hr prior to semen collection. Semen was collected by masturbation into sterile 50 mL sample cups and incubated for a minimum of 15 min post-ejaculation at RT to allow liquefaction. Semen volume was measured, and the density gradient sperm separation procedure (Irvine Scientific) was used to extract motile sperm from liquefied ejaculates. Briefly, 1.5 mL of liquified semen was carefully layered over 1.5 mL of Isolate^®^ (90% density gradient medium, Irvine Scientific) at RT, and centrifuged at 300 g for 20 min. Following centrifugation, the upper layer containing dead cells and seminal plasma was carefully removed without disturbing the motile sperm pellet in the lower layer. The sperm pellet was then washed twice with the sperm washing medium (Irvine Scientific) by centrifugation at 300 g for 10 min. Finally, the purified motile sperm pellet was resuspended in the sperm washing medium, and an aliquot was taken for determination of sperm count and motility using CASA. All semen samples used in the functional assays exceeded lower reference limits for sperm count (15 × 10^6^ total sperm mL^-1^) and total motility (40%) as indicated by WHO guidelines.^[51]^

### 4.7. Sperm count and motility using CASA

The Hamilton-Thorne computer-assisted sperm analyzer, 12.3 version, was used for the sperm count and motility analysis in all experiments unless stated otherwise. This device consists of a phase-contrast microscope (Olympus CX41), a camera, an image digitizer, and a computer with Hamilton-Thorne Ceros 12.3 software to save and analyze the acquired data. For each analysis, 4.4 μL of the semen sample was inserted into MicroTool counting chamber slides (Cytonix). Then, six randomly selected microscopic fields, near the center of the slide, were imaged and analyzed for progressively motile and non-progressively motile sperm count. The parameters that were assessed by CASA for motility analysis were as follows: average pathway velocity (VAP: the average velocity of a smoothed cell path in μm s^-1^), the straightline velocity (VSL: the average velocity measured in a straight line from the beginning to the end of the track in μm s^-1^), the curvilinear velocity (VCL: the average velocity measured over the actual point-to-point track of the cell in μm s^-1^), the lateral head amplitude (ALH: amplitude of lateral head displacement in μm), the beat cross-frequency (BCF: frequency of sperm head crossing the sperm average path in Hz), the straightness (STR: the average value of the ratio VSL/VAP in %), and the linearity (LIN: the average value of the ratio VSL/VCL in %). PM sperm were defined as having a minimum of 25 μm s^-1^ VAP and 80% STR.^[66]^ The complete parameters of the Hamilton-Thorne Ceros 12.3 software are listed in **Table S2**.

### 4.8. Sperm escape assay

This assay was conducted using whole semen and purified motile sperm at the starting concentration of 10 x 10^6^ PM sperm mL^-1^. Briefly, 40 μL aliquots of purified motile sperm or whole semen were transferred to individual 0.2 mL PCR tubes. Sperm count and motility were performed again on each 40 μL aliquot using CASA. This count serves as the original (untreated) concentration of sperm for evaluating the agglutination potencies of respective Ab constructs. Following CASA, 30 μL of purified motile sperm or native semen was added to 0.2 mL PCR tubes containing 30 μL of Ab constructs, and gently mixed by pipetting. The tubes were then held fixed at 45 ° angles in a custom 3D printed tube holder for 5 min at RT. Following this incubation period, 4.4 μL was pipetted from the top layer of the mixture with minimal perturbation of the tube and transferred to the CASA instrument to quantify the number of PM sperm. The percentage of the PM sperm that escaped agglutination was computed by dividing the sperm count obtained after treatment with Ab constructs by the original (untreated) sperm count in each respective tub followed by multiplication with 2 to correct for the 2-fold dilution that occurs upon Ab-treatment. Each experimental condition was evaluated in duplicates on each semen specimen, and the average from the two experiments was used in the analysis. At least 6 independent experiments were done with at least 6 unique semen samples.

### 4.9. Agglutination kinetics assay

#### 4.9.1. FIF-N-Film vs IgG-N-Film

This assay was conducted using both whole semen and purified motile sperm at the starting concentration of 2 x 10^6^ PM sperm mL^-1^, 10 x 10^6^ PM sperm mL^-1^, and 50 x 10^6^ PM sperm mL^-1^. Briefly, 4.4 μL of purified motile sperm or whole semen was added to 4.4 μL of Ab constructs in 0.2 mL PCR tubes, and mixed by gently pipetting up and down three times over 3 s. A timer was started immediately while 4.4 μL of the mixture was transferred to chamber slides with a depth of 20 μm (Cytonix), and video microscopy (Olympus CKX41) using a 10x objective lens focused on the center of the chamber slide was captured up to 90 s at 60 frames s^-1^. PM sperm count was measured by CASA every 30 s up to 90 s. The reduction in the percentage of the PM sperm at each time point was computed by normalizing the PM sperm count obtained after Ab-treatment to the PM sperm count obtained after treatment with sperm washing medium. Each experimental condition, except for 50 x 10^6^ PM sperm mL^-1^, was evaluated in duplicates on each semen specimen, and the average from the two experiments was used in the analysis. At least 6 independent experiments were done with at least 6 unique semen samples.

#### 4.9.2. FIF-N vs FIF-Expi293

This experiment was conducted using the starting concentration of 10 x 10^6^ PM sperm mL^-1^. Briefly, 4.4 μL of purified motile sperm was added to 4.4 μL of Ab constructs in 0.2 mL PCR tubes, and mixed by gently pipetting up and down three times over 3 s. A timer was started immediately while 4.4 μL of the mixture was transferred to chamber slides with a depth of 20 μm (Cytonix), and video microscopy (Olympus CKX41) using a 10x objective lens focused on the center of the chamber slide was captured up to 90 s at 60 frames s^-1^. PM sperm count was measured by CASA every 30 s up to 90 s. The reduction in the percentage of the PM sperm at each time point was computed by normalizing the PM sperm count obtained after Ab-treatment to the PM sperm count obtained after treatment with sperm washing medium. At least 3 independent experiments were done with at least 3 unique semen samples.

### 4.10. CVM collection and processing

CVM was collected as previously described.^[26]^ Briefly, undiluted CVM secretions, averaging 0.5 g per sample, were obtained from women of reproductive age, ranging from 20 to 44 years old, by using a self-sampling menstrual collection device (Instead Softcup). Participants inserted the device into the vagina for at least 30 s, removed it, and placed it into a 50 mL centrifuge tube. Samples were centrifuged at 230 g for 5 min to collect the secretions. Samples were collected at various times throughout the menstrual cycle, and the cycle phase was estimated based on the last menstrual period date normalized to a 28-day cycle. Samples that were non-uniform in color or consistency were discarded. Donors stated they had not used vaginal products nor participated in unprotected intercourse within 3 days before donating. All samples had pH < 4.5.

### 4.11. Fluorescent labeling of purified sperm

Purified motile sperm were fluorescently labeled using Live/Dead Sperm Viability Kit (Invitrogen Molecular Probes), which stains live sperm with SYBR 14 dye, a membrane-permeant nucleic acid stain, and dead sperm with propidium iodide (PI), a membrane impermeant nucleic acid stain. Briefly, SYBR 14 stock solution was diluted 50-fold in sperm washing media. Next, diluted SYBR 14 and PI dye (5 μL) were added to purified sperm (1 mL) resulting in final SYBR 14 and PI concentration of 200 mm and 12 μm respectively. The sperm-dye solution was incubated for 10 min at 36 °C. Then, the solution was washed twice using the sperm washing medium to remove unbound fluorophores by centrifuging at 300 g for 10 min. Next, the labeled motile sperm pellet was resuspended in the sperm washing medium, and an aliquot was taken for determination of sperm count and motility using CASA.

### 4.12. Multiple particle tracking studies

To mimic the dilution and neutralization of CVM by alkaline seminal fluid, CVM was first diluted three-fold using sperm washing medium and titrated to pH 6.8-7.1 using small volumes of 3 N NaOH. The pH was confirmed using pH test strips. Next, Ab constructs or anti-RSV IgG1 control (4 μL) was added to diluted and pH-adjusted CVM (60 μL) and mixed well in a CultureWell™ chamber slide (Invitrogen) followed by the addition of 1 x 10^6^ PM sperm mL^-1^ of fluorescently labeled sperm (4 μL). Once mixed, sperm, Ab, and CVM were incubated for 5 min at RT. Then, translational motions of the sperm were recorded using an electron-multiplying charge-coupled-device camera (Evolve 512; Photometrics, Tucson, AZ) mounted on an inverted epifluorescence microscope (AxioObserver D1; Zeiss) equipped with an Alpha Plan-Apo 20/0.4 objective, environmental (temperature and CO2) control chamber, and light-emitting diode (LED) light source (Lumencor Light Engine DAPI/GFP/543/623/690). 15 videos (512□×□512 pixels, 16-bit image depth) were captured for each Ab condition with MetaMorph imaging software (Molecular Devices) at a temporal resolution of 66.7□ms and spatial resolution of 50 nm (nominal pixel resolution, 0.78 μm per pixel) for 10 s. Next, the acquired videos were analyzed via a neural network tracking software modified with standard sperm motility parameters (**Table S2**) to determine the percentage of PM sperm.^[54]^ At least 6 independent experiments were performed, each using a unique combination of CVM and semen specimens.

### 4.13. In vivo surrogate efficacy studies

On the test day, each sheep received a randomized unique Ab treatment and all sheep were dosed with the same semen mixture that was pooled from 3-5 donors. Briefly, placebo film or FIF-N-film (provided under blind to the animal facility) or saline were instilled into sheep’s vagina and incubated for 30 mins, followed by thorough mixing using a vaginal dilator for 15 strokes. Next, 1 mL of pooled whole semen was pipetted into the sheep’s vagina, followed by simulated intercourse with a vaginal dilator for 5 strokes. Two minutes after the introduction of semen, fluids from the sheep vagina were recovered and assessed for the PM sperm count in a hemocytometer (Bright-Line™ Hemacytometer) under a light microscope (Olympus IX71) using a 20X objective with Thorlabs camera. Each Ab condition was repeated three more times in the same group of sheep (n = 5) with at least 7 days interval in between experiments. Treatments and quantifications were performed in a blinded fashion.

### 4.14. Statistical analysis

All analyses were performed using GraphPad Prism 8 software. For multiple group comparisons, P values were calculated using a one-way ANOVA with Dunnett’s multiple comparisons tests. To compare the percent reduction of PM sperm in vitro by IgG-N-Film vs FIF-N-Film, using whole semen as well as purified semen, one-tailed t-test was performed. Similarly, the comparison between control- and anti-sperm Ab-treated fluorescent PM sperm was performed using a one-tailed t-test. Lastly, to compare the percent reduction of PM sperm in vivo by Placebo-Film vs FIF-N-Film one-tailed t-test was performed. In all analyses, α = 0.05 for statistical significance. All data are presented as the mean ± standard deviation.

## Acknowledgements

We thank Dr. Deborah O’Brien for providing the CASA instrument and her assistance in setting up the CASA measurements. This work was financially supported by the Eshelman Institute of Innovation (S.K.L.); The David and Lucile Packard Foundation (2013-39274; S.K.L); National Institutes of Health under grants R56HD095629 (S.K.L.), U54HD096957 (T.R.M. and S.K.L.), R43HD094454 (T.R.M.) and R44HD097063 (T.R.M.); National Science Foundation (DMR-1810168; S.K.L.); and PhRMA Foundation Graduate Fellowship (B.S.).

## Table of Content

There is a considerable unmet need for safe non-hormonal on-demand contraception. In this work, a hexavalent sperm-binding antibody termed as “Fab-IgG-Fab” (FIF) is produced using a cGMP-compliant system and formulated into a water-soluble vaginal film using polyvinyl alcohol. FIF-Film effectively eliminates 100% of progressively motile sperm in sheep’s vagina within 2 minutes via agglutination and Fc-mediated muco-trapping.

Bhawana Shrestha, Kathleen Vincent, Alison Schaefer, Yong Zhu, Gracie Vargas, Massoud Motamedi, Kelsi Swope, Josh Morton, Carrie Simpson, Henry Pham, Miles B. Brennan, Michael H. Pauly, Larry Zeitlin, Barry Bratcher, Kevin J. Whaley, Thomas R. Moench, Samuel K. Lai*

---

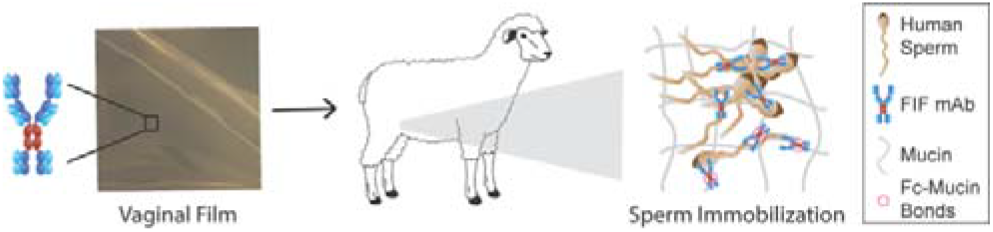
Hexavalent Sperm-Binding IgG Antibody Released from Self-Dissolving Vaginal Film Enables Potent, On-Demand Non-Hormonal Female Contraception.

