## Supplementary File for "Hexavalent Sperm-Binding IgG Antibody Released from Self-Dissolving Vaginal Film Enables Potent, On-Demand Non-Hormonal Female Contraception"

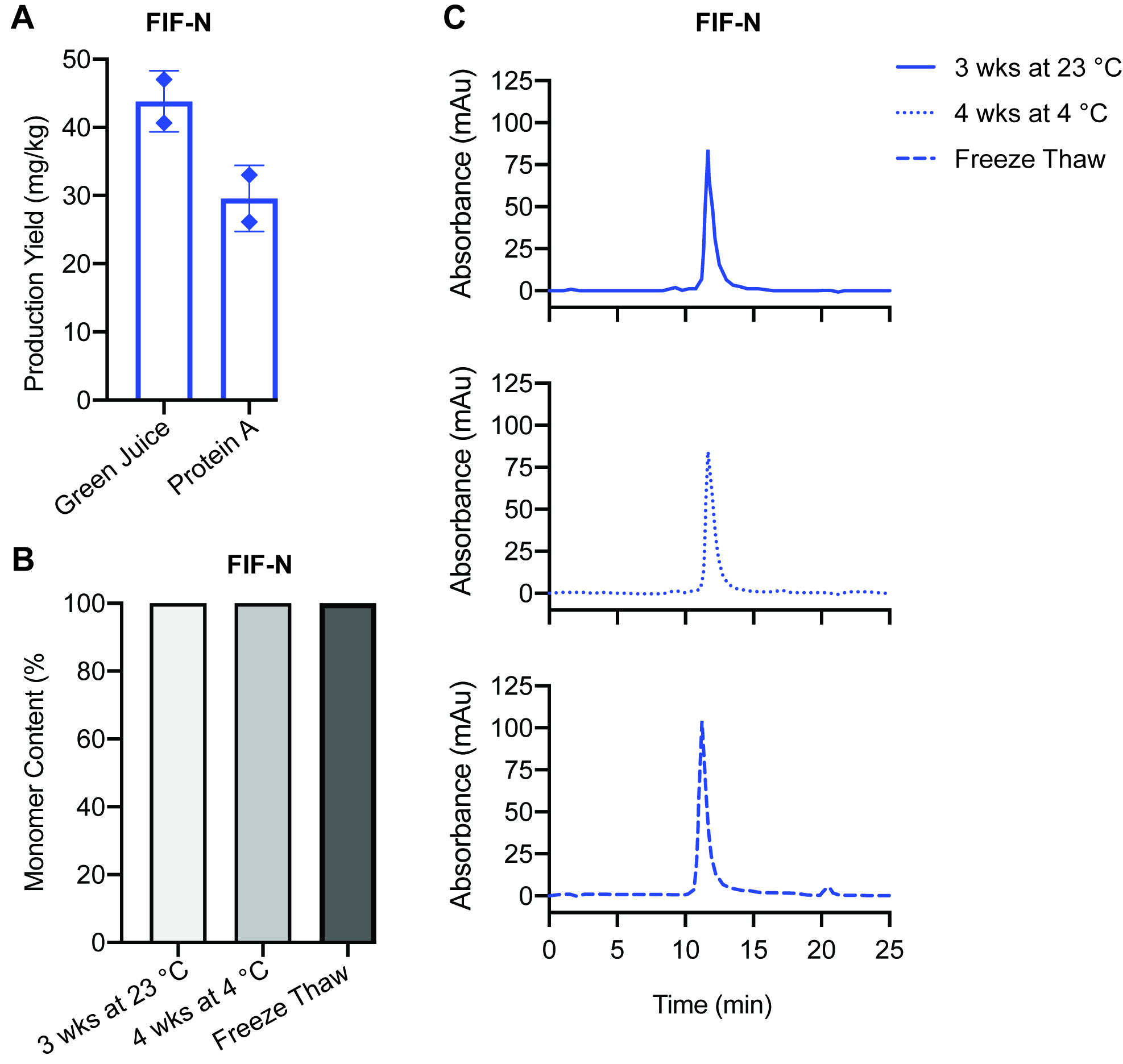

**Figure S1. Additional characterization of *Nicotiana*-produced FIF. (A)** The production yield of IgG and FIF from *Nicotiana* *benthamiana* expression (Green Juice) followed by purification using protein A chromatography. Data were obtained from 2 independent transfections. Lines indicate arithmetic mean values and standard deviation. **(B)** Demonstration of the homogeneity of the FIF-N under different storage conditions using size exclusion chromatography (SEC) analysis. Y-axis indicates the total percentage of Abs representing their theoretical molecular weights. **(C)** SEC curves of the FIF-N stored under different conditions.

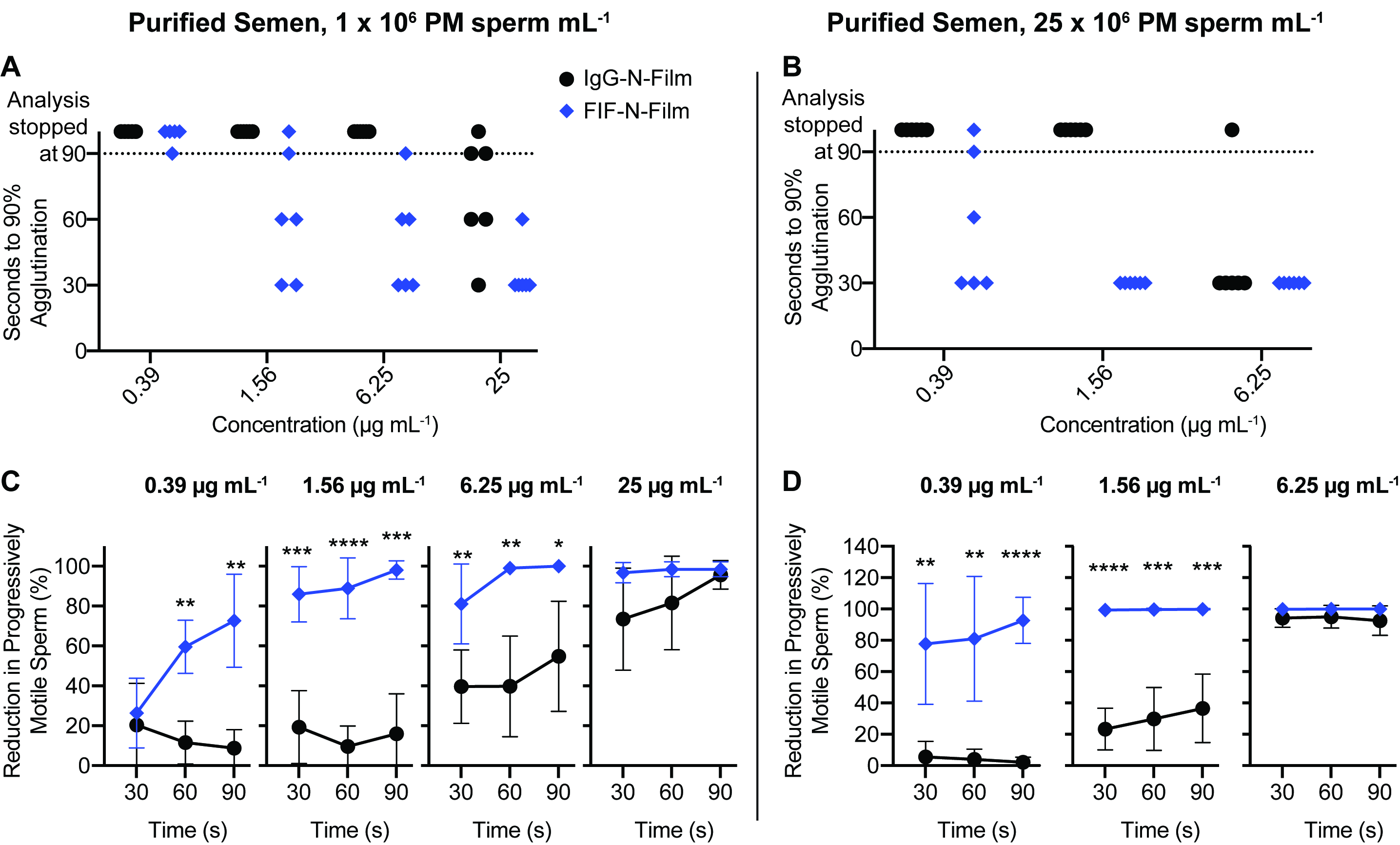

**Figure S2. FIF-N-Film demonstrates faster agglutination kinetics than IgG-N-Film at both low and high sperm concentration. (A)** Sperm agglutination kinetics of the IgG-N-Film and FIF-N-Film measured by quantifying time required to achieve 90% agglutination of PM sperm compared to media control using a final concentration of 1 x 10^6^ PM sperm mL^-1^ and **(B)** 25 x 10^6^ PM sperm mL^-1^. **(C)** The rate of sperm agglutination of the IgG-N-Film and FIF-N-Film measured by measuring the reduction in the percentage of PM sperm at three different time points after Ab-treatment compared to negative control using a final concentration of 1 x 10^6^ PM sperm mL^-1^ and **(D)** 25 x 10^6^ PM sperm mL^-1^. Data were obtained from N = 6 independent experiments with 6 different semen donors. The experiment involving 1 x 10^6^ PM sperm mL^-1^ was performed in duplicates and averaged. P values were calculated using a one-tailed t-test. *P < 0.05, **P < 0.01, ***P < 0.001 and ****P < 0.0001. Data represent mean ± standard deviation.

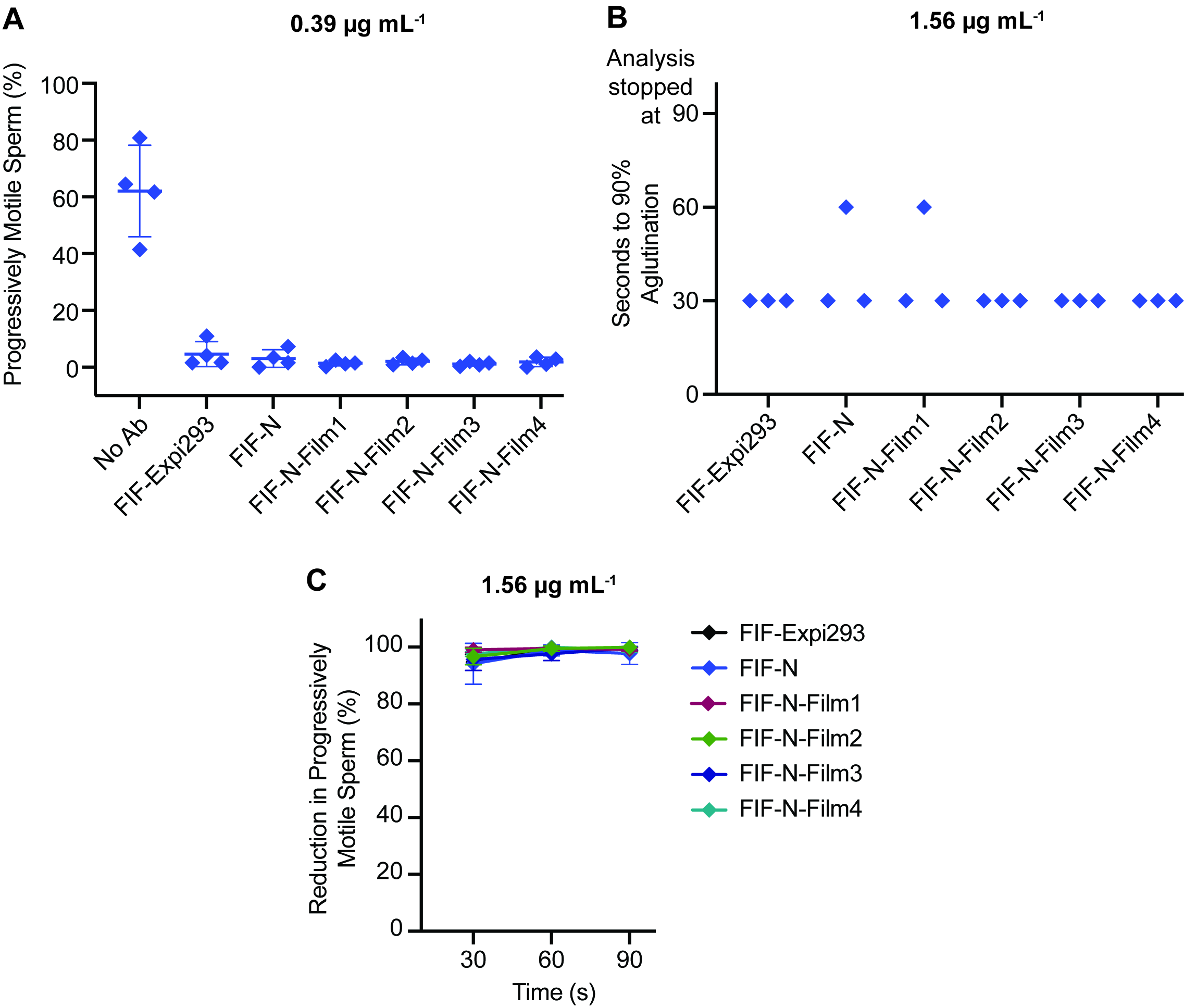

**Figure S3. *Nicotiana*-produced FIF exhibit agglutination comparable to Expi293-produced FIF. (A)** Sperm agglutination potency of the indicated mAbs determined by quantifying PM sperm that escaped agglutination after Ab-treatment compared to pre-treatment condition using CASA. FIF-N-Film1 and FIF-N-Film2 are the film duplicates from one batch whereas FIF-N-Film3 and FIF-N-Film4 are the duplicates from a separate batch. **(B)** Sperm agglutination kinetics of the indicated mAbs measured by quantifying the time required to achieve 90% agglutination of PM sperm compared to media control. **(C)** The rate of sperm agglutination of the indicated mAbs measured by measuring the reduction in the percentage of PM sperm at three different time points after Ab-treatment compared to the negative control. Purified sperm at the final concentration of 5 x 10^6^ PM sperm mL^-1^ was used for all experiments. A significant difference was not observed between the agglutination potency and kinetics of Expi293- and *Nicotiana*- produced mAbs upon one-way ANOVA with Dunnett’s multiple comparisons tests. Agglutination potency data were obtained from N = 4 independent experiments with 3 unique semen specimens. Agglutination kinetics data were obtained from N = 3 independent experiments with 3 unique semen specimens. Data represent mean ± standard deviation.

**Table S1.** Safety parameter results for IgG-N-Film and FIF-N-Film.

| Test Method | IgG-N-Film | FIF-N-Film |
| --- | --- | --- |
| Endotoxin  Bioburden | Film1: <0.958  Film3: <26.1  Film1: 0  Film3: <1 | Film1: 1.505  Film3: <0.953  Film1: 2  Film3: 0 |

**Table S2.** The sperm motility parameters of the Hamilton-Thorne Ceros 12.3.

| Parameter | Value | Parameter | Value |
| --- | --- | --- | --- |
| Frames Per Sec  No. of Frames  Minimum Cell Size  Default Cell Size  Minimum Contrast  Default Cell Intensity  Chamber Depth | 60  60  3 pixels  6 pixels  80  20  20 μm | Path Velocity (VAP)  Straightness (STR)  VAP Cutoff  VSL Cutoff  Slow Cells  Standard Objective  Magnification | 25 μm s^-1^  80 %  10 μm s^-1^  0 μm s^-1^  Motile  10X  1.87 |
